# Calibrating for absolute microbiome abundances without spike-ins

**DOI:** 10.64898/2026.02.26.708180

**Authors:** Nimrod T. de Wit, Amulya Baral, Alessandro Fuschi, Guusje Jacobs, Sharona de Rijk, Rozemarijn Q. J. van der Plaats, Ágnes Becsei, Jesse Kerkvliet, Rebecca Freitag, Martina Vojtková, Christian Brinch, Heike Schmitt, Patrick Munk

## Abstract

Metagenomics is a widely used approach in microbiome research. However, a major limitation of metagenomic datasets is their compositional nature, which prevents direct quantification of absolute abundances and complicates cross-sample comparisons. Existing strategies for absolute quantification typically require additional experiments or spike-in controls. Here, we introduce the MetaGenome Calibrator (MGCalibrator), a new tool that enables spike-in free, absolute abundance estimation based on routine DNA concentration measurements.

We validated the accuracy of absolute abundances obtained with MGCalibrator against qPCR for 5 targets. Our results show a strong correlation with qPCR data, indicating that MGCalibrator enables qPCR-like trend analyses. For *Bacteroides dorei*, the estimated abundances were highly similar between the two methods (r2 = 0.98, y = 1.00x). For other targets like crAssphage or the bacterial 16S rRNA gene, qPCR values were underrepresented by a factor of 7 or overrepresented by a factor of 4. Benchmarking with synthetic microbiome data demonstrated that our method accurately determines copy numbers in sequencing datasets, and application to whole-cell mock community samples produced expected values based on known extraction biases. In an extraction-bias-free experiment, MGCalibrator accurately quantified genome copy numbers within a twofold range in 98% of cases and determined 16S rRNA gene copies within 1.6-fold or less.

Finally, we applied MGCalibrator to track temporal trends in antibiotic resistance genes (ARGs) in wastewater treatment plants in two Dutch provincial capitals. We observed an overall increase in ARGs—such as *sul2* in Utrecht and *qnrS5* in Houtrust—likely driven by rising bacterial loads. Our findings demonstrate that MGCalibrator provides robust calibration of metagenomic data, paving the way for metagenomics to play a central role in future surveillance by enabling trend analysis across thousands of genetic targets, similar to the capabilities of qPCR for individual genes. The source code and documentation for MGCalibrator are available at github.com/NimroddeWit/MGCalibrator.

## Introduction

Microbial communities play central roles in human health^1^, environmental processes^2^, and engineered systems^3^, making accurate characterization of these communities essential for understanding their dynamics and function. Many biologically relevant phenomena, such as pathogen overgrowth, dysbioses, bloom-collapse events^4^ in metagenomic samples depend not only on the presence of specific taxa but also their absolute abundances^5,6^. Yet most microbiome studies rely on metagenomic sequencing, which reports only relative abundances.

Because a metagenome is a sequencing-dependent subsample of the true DNA pool, read counts cannot be directly compared across samples, and the overall scale of each sample, or the microbial load cannot be recovered from metagenomic sequencing alone. This difference between absolute and compositional data can significantly distort biological interpretation and generate misleading conclusions.^7–10^

Previously, multiple strategies have been proposed to recover or estimate the missing scale of 16S rRNA gene amplicon sequencing or shotgun metagenomic data. PCR-based methods, including quantitative PCR (qPCR) and digital PCR (dPCR) can provide precise measurements of specific genetic targets^8,11^, but require highly specific primers. This is especially challenging in complex sample matrices.

Another strategy involves adding genetic material of known quantity (spike-ins), such as whole cells^12^ or purified nucleic acids of a known taxon, or synthetic standards^12,13^. Although spike-ins provide a verifiable quantity of reference genetic material, they require additional preparation steps, increase sequencing costs and are not possible to apply post-hoc to the hundreds of thousands of sequenced metagenomes. Moreover, since the spiked organisms should be absent from the metagenomic sample, prior knowledge of the sample content is required.

More recently, progress has been made toward *in silico* estimation of the missing scale. For example, a recent study expanded on the statistical functionality of ALDEX2, by incorporating this scale uncertainty into the differential abundance estimating model^14^. Such an approach allows the user to account for relative differences in scale based on additional outside measurements like flow cytometric cell counting.

We sought to provide and validate a metagenomic data-based quantification workflow that leverages concentration measurements to enable spike-in free absolute abundance estimation. Our proposed workflow does not require additional laboratory techniques but instead relies on DNA measurements routinely undertaken prior to sequencing. Although this concept was previously proposed by Jiang *et al*.^15^, their study was not published, nor experimentally validated. Yin *et al*.^16^ applied a similar approach but likewise lacked experimental confirmation. To this end, we introduce a user-friendly command-line tool called MGCalibrator, that corrects for incomplete sequencing and accounts for measurement and sampling uncertainties when converting read counts to absolute abundances. We validate our approach by benchmarking against qPCR measurements, through simulations, and by analyzing mock communities. Finally, we attempt time-series analyses of AMR genes in wastewater, laying the foundation for metagenomic spike-less absolute quantification applicable to both past and future studies.

## Results

### Framework for absolute microbiome abundance estimation

We developed a metagenomic calibration tool named MGCalibrator, which enables absolute quantification and uncertainty estimation, based on just extracted DNA concentration and metagenomic read classifications (Figure 1).

**Figure 1.**
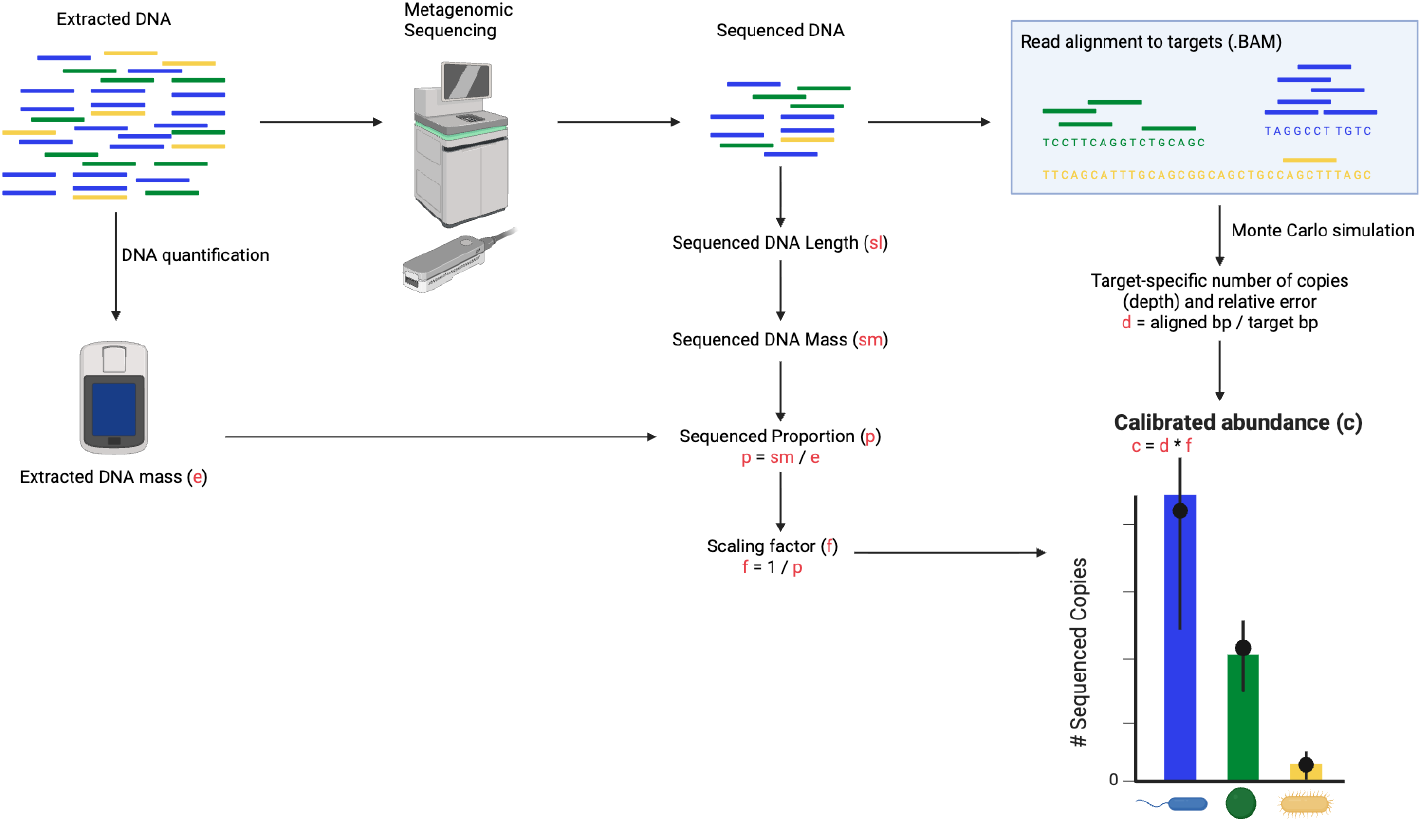
Graphical overview of MGCalibrator. Extracted DNA is quantified to obtain the total extracted mass (e). Following metagenomic sequencing, the total sequenced DNA length (sl) is converted to sequenced DNA mass (sm), allowing calculation of the sequenced proportion (p = sm / e). A scaling factor (f = 1 / p) is derived to correct for subsampling during sequencing. Reads are aligned to target sequences (BAM), and target-specific depth (d = aligned bp / target bp) is estimated using Monte Carlo simulation to account for relative error. Calibrated abundance (c) is then computed as c = d × f, yielding target-specific copy number estimates with associated uncertainty.

Because the mass of DNA is known (650 dalton/bp), we can calculate what all the sequenced ssDNA molecules weighed in total. Comparing this value to the mass of extracted DNA, we can calculate the fraction that was actually sequenced. The reciprocal of this fraction serves as a mass-calibration scaling factor. When this scaling factor is multiplied with sample-wise abundances, the resulting values represent DNA mass-calibrated abundance, which allow absolute inter-sample comparisons.

In this framework, samples with low sequencing throughput or high extracted DNA yield, have low sequenced fraction and therefore receive a large scaling factor.

For example, consider a sample from which we extract 1 µg of DNA and sequence 50 million paired-end reads of 150 bp, you would end up sequencing just 15 Gbp weighing 16.2 pg, corresponding to a 62000th of the total DNA, giving us the scaling factor. All read-derived quantities can therefore be multiplied with this scaling factor when estimating their absolute abundances. Under this scaling, an *E. coli* with a depth of coverage (hereafter referred to as depth) of 20x in the sequencing data corresponds to approximately 1.2 million genome copies in the DNA extract.

However, a depth value alone does not give any information on certainty, even though certainty can differ significantly. The number of reads mapped to each reference is subject to stochastic variation. If the same library were sequenced again, slightly different read counts would be observed for each reference due to this random sampling. This variation is increased by the fact that the same mean depth value can correspond to vastly different numbers of reads, depending on the length of the reference sequence. A mean depth of 1x across the human genome corresponds to millions of short reads, whereas a depth of 1x across a single gene might correspond to only ∼10 reads. Thus, for identical depth values, the proportional error is substantially larger for shorter reference sequences.

To account for this variability, our approach simulates the variation in the number of reads mapping to the targets of choice using Poisson statistics and Monte Carlo simulation. Therefore MGCalibratortakes as input a) reads aligned to target sequences across one or more samples and b) the mass of extracted DNA for those samples. Finally, MGCalibrator returns the number of reference molecules of interest with confidence intervals (see Methods).

### Validation of calibration with qPCR

To test our approach, we took 40 samples from 18 wastewater treatment plants (WWTPs) across the Netherlands and analysed them with both metagenomics and qPCR. We validated MGCalibrator output against qPCR measures for five targets: the human gut commensal *Bacteroides dorei*, the human fecal indicator crAssphage, the bacterial 16S rRNA gene, and two low-prevalence antibiotic resistance genes (ARGs): *vanA* (vancomycin resistance) and *bla*_CTX-M_ (extended-spectrum β-lactamase) (Figure 2).

**Figure 2.**
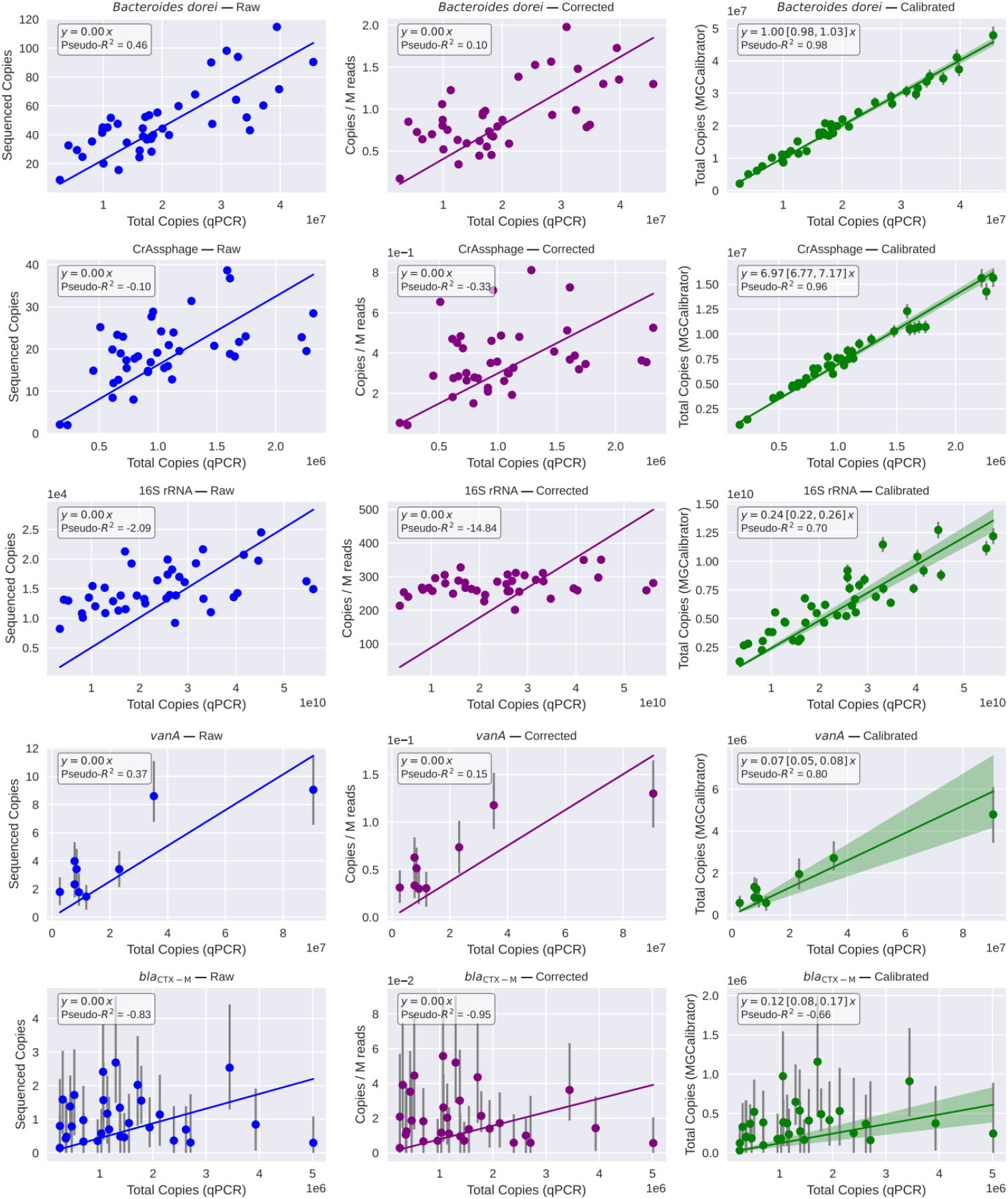
Comparison of abundances estimated with qPCR and metagenomics. For each metagenome, five targets were quantified using both qPCR and metagenomics. Left column: y axes are sequencing depth over the reference. Middle column: y axes represent depth corrected for millions of reads. Right column: y axes represent the DNA mass-calibrated gene/genome counts, i.e., the total number of copies in the DNA extract. For each target, only samples where both instruments detected signals were included. Error bars represent 95% confidence intervals estimated using Monte Carlo simulations. Lines show weighted least-squares linear fits (inverse-variance weighted, forced through the origin); shaded bands and slope intervals denote 95% model-based confidence intervals.

The correlation coefficient drastically improved after calibration. The correlation over the sample set was especially high for our tested species, CrAssphage (0.96) and *B. dorei* bacterial genome (0.98). It was lower for the combined 16S rRNA genes (0.70) and *vanA* (0.82). In none of the metagenomes, we sequenced *bla*_CTX-M_ to a 3+ depth, giving it a large confidence interval and low correlation to qPCR (Figure 2).

Among the 14 samples with a positive qPCR result, *vanA* was detected by shotgun metagenomics only in the 9 samples with the highest concentrations. All samples with concentrations ≤ 2.55 × 10^4^ copies/µL were negative for *vanA* by metagenomics. In contrast, for *bla*_CTX-M_ there was no connection between qPCR concentration and detection by metagenomics, as both the sample with the highest concentration (5.01 × 10^4^ copies/µL) and the sample with the lowest concentration (2.51 × 10^3^ copies/µL) were negative by metagenomics.

Except for *B. dorei*, all of the slopes were very different from the expected y = 1x, being 6.97 for crAssphage, 0.24 for 16S rRNA, and 0.07 for *vanA*. For *B. dorei* we used species-specific 16S rRNA gene primers to quantify it. This species has 7 16S rRNA copies. When correcting for this, qPCR and metagenomics are in almost perfect agreement (slope = 1.00). The pronounced underestimation of crAssphage by qPCR, yielding approximately seven-fold fewer copies than metagenomic quantification, cannot be explained at this time. Inspecting the read depths over the length of the genome, did not reveal any gross errors, and the primed genome area was indeed conserved, with a very low rate of read disagreement (Supplementary Figure 1). Additionally, depth at the primed sites did not differ significantly from the depth at other sites, as confirmed by bootstrapping (Supplementary Figure 2). For 16S rRNA and *vanA*, metagenomics detected approximately one-quarter and one-seventeenth of the reads, respectively, compared with the qPCR quantities. The qPCR primers of both targets may be less specific, potentially leading to an overestimation of abundance. However, for *vanA* the concentrations were close to the metagenomic limit of quantification, which might also be a factor contributing to the underestimation.

### Validation using synthetic and mock microbiome data

Unspecific mapping, deletions and insertion events can all contribute to differences between real composition and a metagenomic abundance estimate. To validate our depth calculation method, we chose to create three *in silico* datasets, varying in sequencing depth. Each dataset consists of thirty species, including plasmids, with sequences varying in length and abundances (Supplementary Table 1). We see that for the majority of sequences the calculated abundance agrees with the synthetic ground truth (Figure 3a). As expected, we found that small sequences indeed have a larger proportional error.

**Figure 3.**
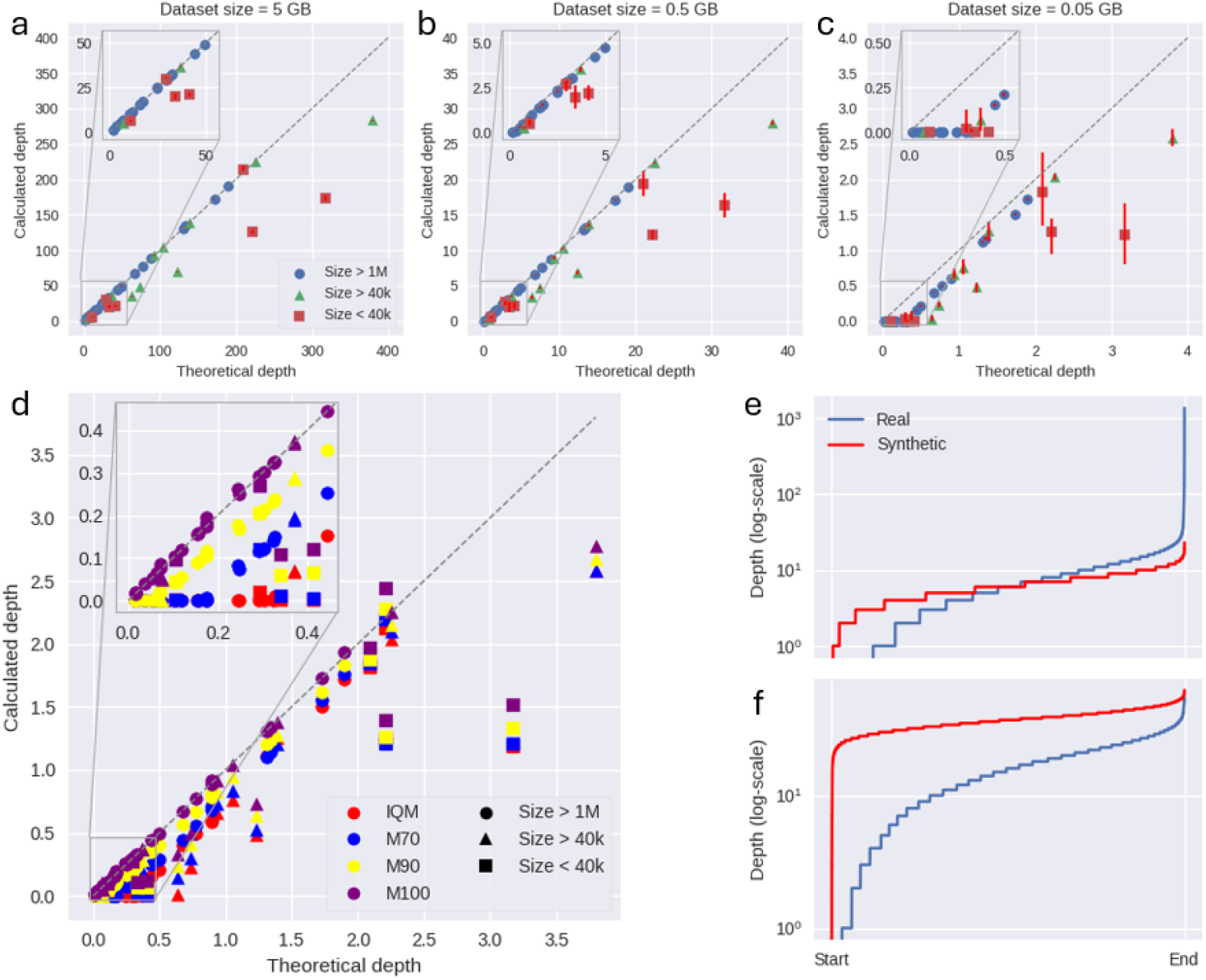
Validation of MGCalibrator using synthetic microbiome data. (a-c) Effect of reference length and sequencing depth on abundance estimation accuracy. Each point represents a sequence in the synthetic dataset (Supplementary Table 1). (d) Effect of depth-calculation method on abundance-estimation accuracy. IQM denotes the interquartile mean of sequence depth within one sequence, while M70, M90, and M100 represent the mean of the central 70%, 90%, and 100% of the sorted depth values, respectively. Based on a 0.05 GB simulated metagenome. (e-f) Depth-of-coverage values for bacterial genomes (e) and crAssphage genomes (f) in a real and a synthetic sample. Depth values were sorted and plotted on a logarithmic scale. The bacterial genome in the real sample is Bacteroides dorei (genome size 5.6 Mb; IQM depth 6.2), whereas the bacterial genome in the synthetic sample is Halovivax ruber (genome size 3.2 Mb; theoretical depth 6.4). For crAssphage (genome size 97 kb), the IQM depth in the real sample is 15.5, and the theoretical depth in the synthetic sample is 37. “Start” and “End” indicate the beginning and end of the genome, respectively.

Depth underestimation observed in smaller reference sequences reflects the presence of non-unique sequences, such as genes or genomic regions shared across elements, where reads can map equally well to multiple references. The way Minimap2 by default resolves ambiguous mappings is by arbitrarily selecting one of the possible references, rather than proportionally distributing reads across identical reference sequences. Consequently, measured depths differ from theoretical expectations and this effect is stronger for plasmids that tend to have higher proportions of shared sequence.

To determine the best way to estimate depth, we tested several methods, varying in the percentage of depth values included in the average calculation. The interquartile mean (IQM) includes 50% middle of the sorted depth values, M70 includes 70%, and so on. Our simulated dataset shows that for M100, the depth most closely matches the ground truth (Figure 3b). However, we also compared the depth profiles of real and synthetic datasets, to see if the depth calculation strategy should be different for real and synthetic datasets. We did observe different depth profiles, with depth profiles in synthetic data being flatter compared to real data (Figure 3c-d).

Additionally, we sequenced duplicated ZymoBIOMICS Microbial Community Standards and compared the number of spiked genome copies to our calibrated abundances. Some microbes are difficult to lyse, which can cause DNA extraction protocols to bias metagenomic results^17^. Therefore, we did not expect to recover proportional numbers of genome copies across species.

But if our method works, we would never expect to estimate more genome copies for a species than the theoretical max, which assumes an unrealistic 100% recovery with no loss. As expected, absolute recovery was <100% in all cases, and furthermore related to species identity (Figure 4a). Gram-positive bacteria have tough peptidoglycan cell walls and are known to be under-sampled in sequencing results^18^. Consistent with that fact, Gram-negative organisms had calibrated abundances closer to the theoretical maximum.

**Figure 4.**
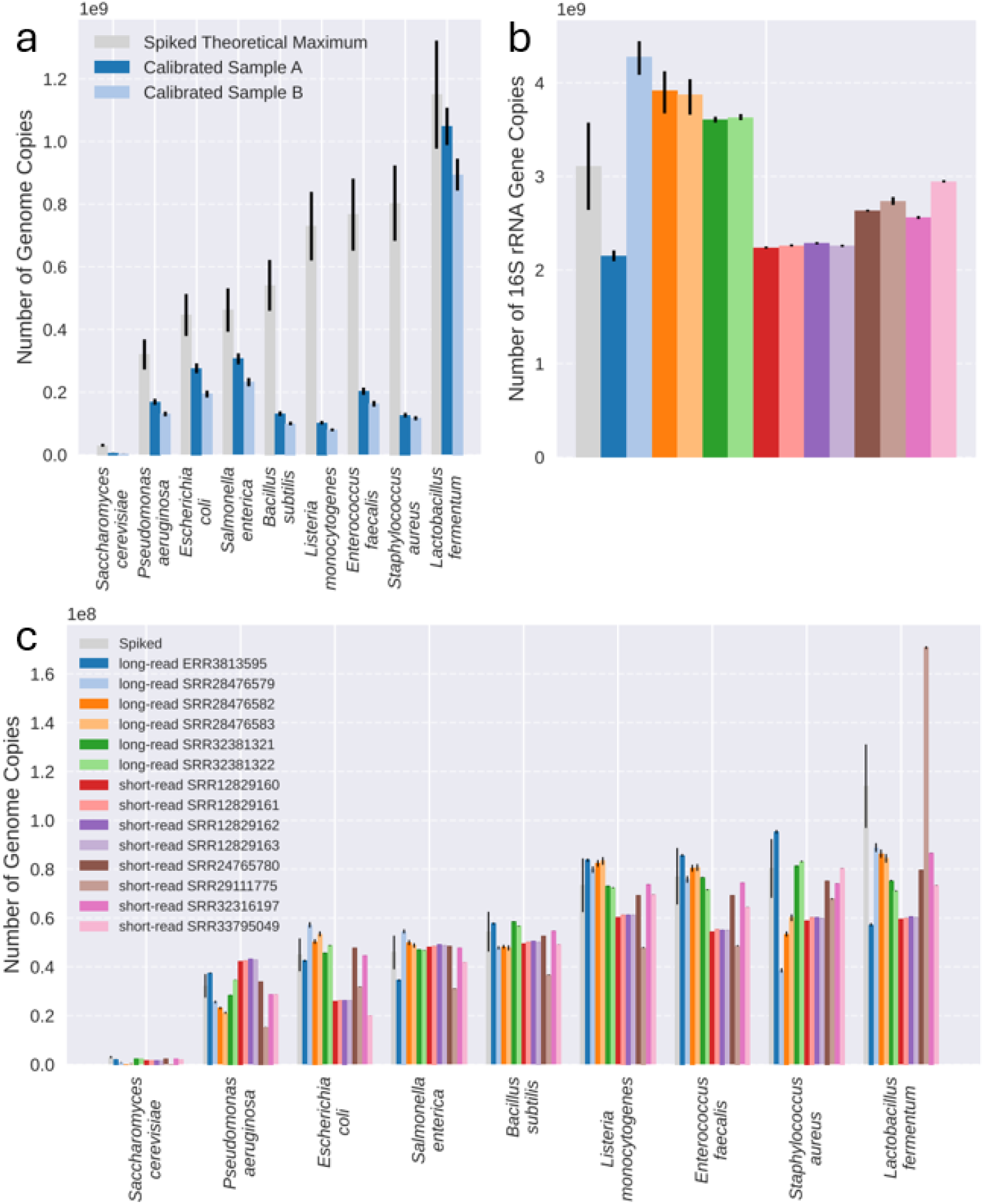
Validation of MGCalibrator using mock microbiome communities. (a) Taxon-specific differential recovery during DNA extraction and sequencing of a whole-cell mock community. Known spike-in quantities (grey) are compared with MGCalibrator-estimated copy numbers for technical replicates A and B. Replicate A showed consistently higher estimated abundances across all spike-in organisms relative to replicate B (two-sided paired t-test, p = 0.012). (b,c) Quantification of 16S rRNA gene copy numbers (b) and species-specific genome copy numbers (c) in six long-read and eight short-read shotgun metagenomic datasets generated from DNA mock communities. Spiked quantities are shown in grey. Across all panels (a–c), error bars for spiked quantities reflect the 15% uncertainty reported by the manufacturer, whereas error bars for MGCalibrator estimates indicate 95% confidence intervals derived from Monte Carlo simulations.

To avoid extraction bias, we applied MGCalibrator to ZymoBIOMICS Microbial Community DNA Standard samples. Short-read (Illumina) and long-read (ONT) shotgun metagenomic datasets were retrieved from the European Nucleotide Archive (ENA), spanning a range of read depths and sequencing platforms (Supplementary Table 2).

Measured copy numbers were largely consistent with theoretical abundances: 57 of 126 measurements fell within the expected error range, and all but four of the remainder deviated by less than twofold (Figure 4c). The four measurements exceeding this threshold all corresponded to *S. cerevisiae* and occurred in the samples with the lowest read depth. This suggests an influence of sequencing depth, consistent with our observations in the synthetic microbiome dataset. Similarly, for these four samples, the number of genome copies in the sequencing dataset was < 1, and M95 substantially mitigated the underestimation observed with IQM, reducing the magnitude of underestimation from 3.8–11,100-fold to 1.4–1.7-fold (Supplementary Figure 3). Among bacteria, *Lactobacillus fermentum* showed the greatest discrepancy.

Because qPCR and MGCalibrator disagreed in the quantification of the 16S rRNA gene, we additionally quantified 16S rRNA in these datasets and compared the results to the theoretical gene copy numbers. Consistent with previous observations, MGCalibrator closely matched the expected values: 6 out of 14 measurements fell within the expected error range, and all remaining samples deviated 1.6-fold or less (Figure 4b).

### ARGs increasing over time in the Netherlands

Because our results showed we could obtain absolute abundances varying like qPCR-values, we set out to perform trend analyses with MGCalibrator. With thousands of known ARGs, it would not be feasible or economical to survey trends of the combined resistome with qPCR, but this could be a prime use for MGCalibrator.

We apply this approach to antibiotic resistance surveillance in wastewater from two provincial capitals in the Netherlands. The surveillance covers a period of 2.5 years, with 7 time points for Utrecht and 8 for Houtrust. We observe that several genes, resistance phenotypes and classes have significantly increased over the past two years. This includes the *qnrS5* fluoroquinolone resistance gene in Houtrust and the *sul2* sulfonamide resistance gene in Utrecht (Figure 5).

**Figure 5.**
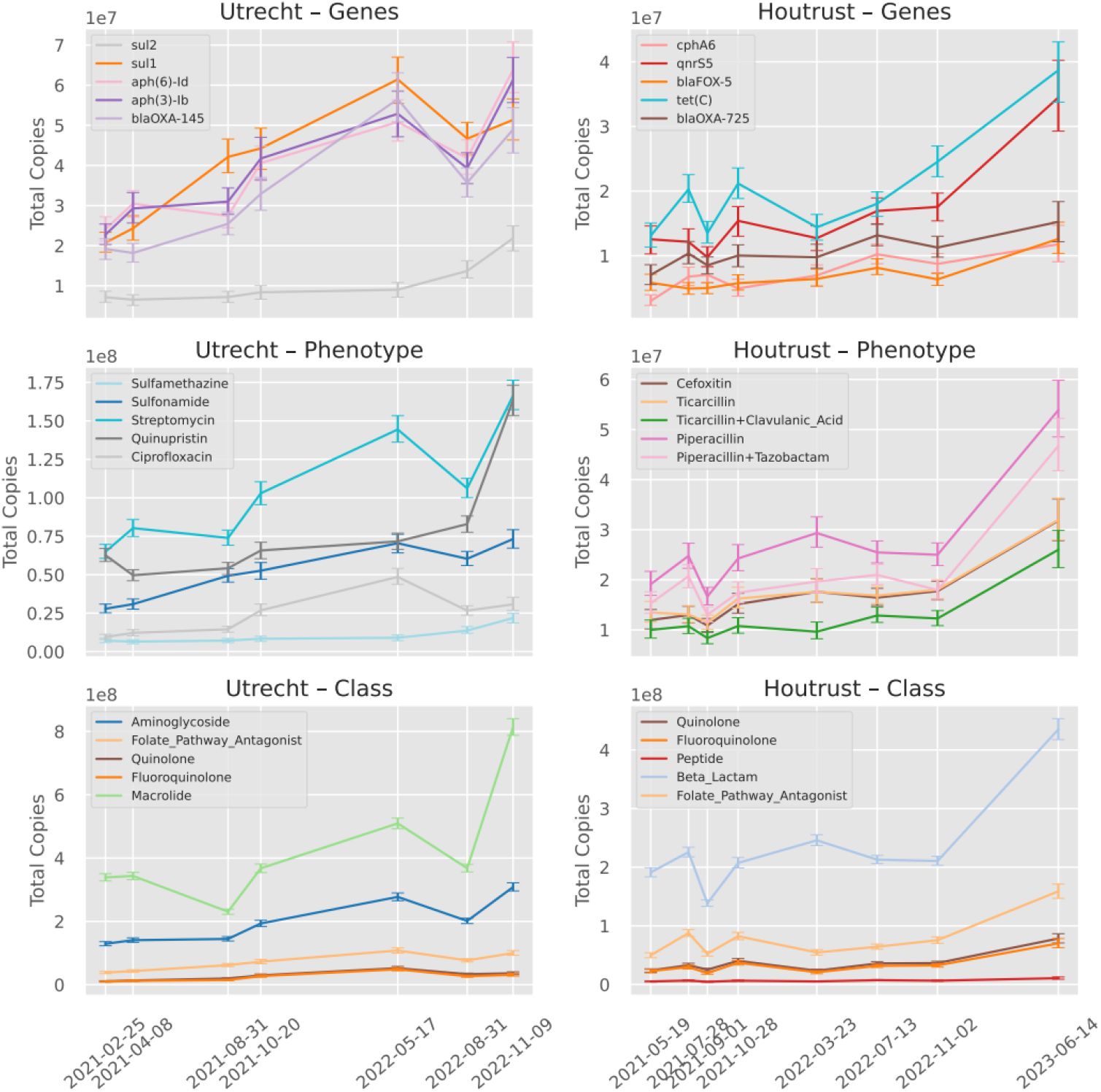
Top five increasing genes, phenotypes, and resistance classes in Utrecht and Houtrust. Wastewater treatment plants in Utrecht and Houtrust were sampled over periods of 1.7 and 2.1 years, yielding 7 and 8 samples, respectively. Phenotype and resistance class abundances were computed by summing the abundances of all annotated genes. Significant trends were identified using Kendall’s τ correlation test, accounting for uncertainty. For clarity, only the five features with the highest Kendall’s τ values are shown. Error bars indicate 95% confidence intervals estimated using Monte Carlo simulations.

In addition, a general increase in antibiotic resistance is observed. Across the two cities, a large proportion of features showed rising abundance, reflected by an overrepresentation of low p-values (Supplementary Figure 4). The observed increase in crAssphage and 16S rRNA genes, combined with a stable ARG/16S rRNA gene ratio, however suggests that this increase in ARGs is rather caused by a higher fecal load late in the sampling campaign (Supplementary Figure 5).

In Utrecht, top rising resistance genes encoded varied phenotypes, to both sulfonamide type drugs, ciprofloxacin and streptomycin. In Houtrust on phenotype level, all the largest increases were associated with beta-lactam resistance phenotypes. *qnr*S5, *sul*1 and other genes known to mobilize especially between Enterobacteriaceae were prominent among rising ARGs.

## Discussion

By integrating the measured DNA mass and the calculated mass of extracted DNA, we were able to estimate absolute abundances akin to qPCR. Although this results in an almost perfect match for *Bacteroides dorei*, there is a discrepancy between sequencing and qPCR for the other targets, with slopes ranging from 0.06 to 6.93. Barlow *et al*. also observed species-specific slopes and suggested it could be caused by a discrepancy in coverage/specificity between the taxon-specific and universal primer sets.^8^ They specifically noticed that *Akkermansia muciniphila* had a 2.5x overrepresentation in sequence-based quantification compared to qPCR, which is similar to our 7x overrepresentation for crAssphage.

Differences observed between qPCR and metagenomic sequencing may not necessarily reflect true biological variation in target abundance, but rather methodological biases inherent to each approach. qPCR quantifies a short, predefined region within a gene and is largely independent of full gene length or overall coverage, but it may underestimate abundance when primer mismatches, subtype diversity, or matrix-associated inhibitors reduce amplification efficiency (Elbait et al.). Metagenomics is generally less susceptible to inhibition effects during detection and quantification, but it is less sensitive than qPCR, relies on random read generation across entire sequences, and often requires a minimum proportion of gene coverage for confident detection (Lagenfeld et al.) as relative error increases with lower coverage depth. Together, these factors can lead to systematic differences in slope or proportional abundance estimates between the two methods, even when the underlying biological signal is similar. We cannot be certain which of the two methods is more accurate, but we do note that MGCalibrated values were all within a factor of 2 of control Zymo DNA standard datasets, while qPCR could disagree with, e.g., 7x for crAssphage. For the 16S rRNA gene, given that MGCalibrator was often close to expected values for ground truth data, but could differ significantly from qPCR abundances, we suggest that qPCR is likely suffering from larger biases. A future study should prioritize including mock communities with known genome copy numbers for species where qPCR primers are available.

Absolute quantification is only useful when the quantity of genomes or genes of interest in the sample exceeds the limit of quantification. Therefore, for low-abundance genes it works poorly (*vanA*) or not at all (*bla*_CTX-M_). However, this is not specific to our method but is inherent to shotgun metagenomics in general. A strength of our approach is that instead of using, for example, entropy-based thresholds (Langenfeld et al.), we opted to include confidence intervals, so even low-abundance entities can result in point estimates with an associated CI. The error estimates provide a direct indication of the reliability of the calibrated abundance estimates. Wide confidence intervals on genes of interest gives the user the chance to prioritize sampling more, i.e., increasing sequencing depth.

MGCalibrator takes a BAM file as input, containing all metagenomic reads aligned to a defined set of reference sequences. We expect MGCalibrator to perform optimally when using alignments to non-redundant reference sets with minimal sequence overlap, thereby reducing ambiguous read mapping. Alternatively, such ambiguity can be mitigated by selecting an appropriate mapping strategy. Ideally, the proportion of reads that map equally well to multiple references should reflect the sequencing depth observed in the unique regions of those references. For gene quantification, ambiguous read mapping is unlikely to pose a problem, given that substantial overlapping regions between distinct genes are rare. An exception would be recombining tetracycline resistance genes like *tet*(/W/32/O)^19^. Additionally, a redundant database containing multiple alleles per gene can be employed, as MGCalibrator has the option to cluster alleles.

Besides using a known reference for alignment, it is also possible to use *de novo* assembled contigs based on the metagenomic reads, though it leads to an increased uncertainty according to Langenfeld et al. So far, MGCalibrator has not been tested on de-novo assembled contigs, but especially for trend analyses in longitudinal surveillance, where our approach should be useful, it would potentially be beneficial to co-assemble metagenomes sharing strains to establish more comprehensive metagenome-assembled genomes databases as previously demonstrated^20^.

To address the issues relating to variable depths of coverage as raised by Elbait et. al^21^, we tested several ways of estimating depth. We observed that M100 (mean of all sequence depth values) performs best as a depth calculation method for synthetic dataset, and M95 outperforms IQM (mean of middle 50% of sorted depth values) for low abundant *S. cerevisiae* in the mock dataset. However, the reason that M95 and M100 work best in these scenarios is likely because the simulated and mock data are a simplified representation of reality, i.e., all genomes providing reads are known. This is not the case with real datasets, as we observed when comparing the depth profiles. In real-world datasets, reads are typically mapped to a subset of the genomes actually present in the sample. This can lead to non-specific alignments, for example to conserved or widely shared housekeeping genes, resulting in localized stacks of reads that are misaligned. Conversely, the reference genomes used for mapping may contain accessory genes that are absent from the genomes in the sample. This mismatch creates gaps or marked reductions in coverage. To avoid including these outlier positions in the depth calculation, we prefer to use the IQM despite the risk that this may lead to a slight underestimation for low abundant organisms. This is therefore the default option of MGCalibrator.

Experiments using the ZymoBIOMICS Microbial Community DNA Standard demonstrate that variability is introduced at multiple stages, including library preparation, sequencing, and downstream bioinformatic analysis. The magnitude of this variability differs between samples and appears to be at least partially dependent on sequencing depth, as the four datasets requiring substantial scaling exhibited markedly reduced detection of *S. cerevisiae*.

Beyond estimating copy numbers in a DNA extract, it is crucial to ensure that the relationship between the original sample and the resulting DNA extract is consistent across all samples. This requires sample preprocessing steps to be standardized, as variability introduced at this stage alters the proportion of the original material that is represented in the extract. Maintaining a fixed and well-defined preprocessing workflow is therefore essential to preserve comparability between samples. For example, in wastewater analysis, processing a defined sample volume by filtration and extracting DNA from the entire filter ensures that each DNA extract represents the same fraction of the original sample. In contrast, centrifugation followed by random subsampling of the pellet will introduce uncontrolled variation in this proportion. Additionally, the capacity of the filter used for DNA extraction should be considered to ensure consistent recovery across samples^8^.

In conclusion, we find that, despite qPCR and metagenomics each showing different target preferences, the linear correlation between the methods can be very strong (R^2^ > 0.96 for *B. dorei* and crAssphage). This means that spikeless absolute quantification allows longitudinal trend analysis, as in the case for thousands of ARGs. It appears that calibrated metagenomics can play a prominent role in future surveillance, due to the fact that it allows trend analysis for millions of ARGs, viruses, other pathogens, as well as toxin genes and other virulence factors. We hope that MGCalibrator will also find use in the re-analysis of novel targets in many unspiked datasets.

## Methods

### Sampling

24-hour flow proportional samples of influent were collected using automated samplers according to the routine sampling regime of the WWTP, at one time-point between July 2020 and June 2023. Sampling was performed by WWTP staff. Samples were kept cool during storage and transportation and filtered soon after; mostly within 36 h after collection. Two times 10 mL influent was filtered on a 0.22 µm filter and this was stored at -20° Celsius. Additionally, two samples containing 375 uL of ZymoBIOMICS Microbial Community Standards (reference number D6300) (∼1.4 × 10^10^ cells/ml, 0.525 × 10^9^ cells per sample) were included for sequencing. See Supplementary Table 3 for a list of all samples, with relevant sequence statistics and accessions.

### DNA extraction and quantification

Filter papers were collected in PowerWater DNA Bead Tubes and fragmented by bead beating using the Vortex Adapter (cat. No. 13000-V1-5/13000-V1-15) at maximal power for 7.5 minutes. Then the protocol of the QIAGEN DNeasy PowerWater kit was followed. DNA quantity of all samples was measured using Qubit® dsDNA HS Assay Kits 1x solution on a Qubit Flex Fluorometer. Qubit measurements were performed according to the manufacturer’s protocol.

### qPCR

All qPCR reactions were conducted in 20 µL, including the reaction mixture as specified in Supplementary Table 4. A total of 5 µL (for 16S rRNA, bla_CTX-M1_, and *vanA*) and 3 µL (for crAssphage and *Bacteroides dorei* (HF183)) DNA template was added to each reaction. All reactions were performed on a LightCycler® 480 Real-Time PCR System (Roche). For primer and probe sequences and concentrations, as well as PCR cycles, see Supplementary Table 4. The annealing temperature was the same for all targets, except for *vanA*, for which the temperature was 58°C.

Synthetic DNA fragments (IDT, US) corresponding to each target gene served as positive controls for generating the standard curves (Supplementary Table 5). Serial dilutions of these fragments were prepared in sheared salmon sperm DNA (5 µg mL^−1^; ThermoFisher, LT) suspended in Tris-EDTA (TE) buffer at pH 8.0 (Sigma Aldrich, Switzerland). All samples were run in technical duplicates. Each PCR plate contained a standard curve consisting of at least five dilution points, also analyzed in duplicate. For each gene, a mean standard curve was produced. Gene concentrations were subsequently determined using this averaged curve.

We evaluated the linear relationship between calibrated metagenomic and qPCR measurements by fitting a line through the origin. We calculated a pseudo-R^2^ as the proportion of variance explained by the linear fit. This value is equivalent to the squared Pearson correlation coefficient between observed and predicted values without an intercept.

### Metagenomics

Library preparation was performed with the Illumina DNA Prep Tagmentation set (reference number 20060059). Illumina DNA/RNA UD Indexes Set D was used, and in one batch of 55 samples, the libraries were multiplexed and sequenced on the NextSeq2000 platform (Illumina), using the NextSeq2000 P4 XLEAP-SBS flowcell (2 × 150-bp paired-end sequencing). This resulted in an average of ∼40 million reads per sample. Sequencing reads were trimmed, filtered, merged, and deduplicated using Fastp^22^ v0.24.0 with default settings. Illumina datasets downloaded from ENA were preprocessed in a similar manner. Nanopore datasets from ENA were processed using Chopper v0.8.0, with parameters `--quality 9 --minlength 500 --headcrop 50 --tailcrop 50;` all other settings were default.

### Obtaining BAM files

BAM files that were used as input to MGCalibrator were obtained in the following manner, with only the set of reference genes or genomes varying between organisms: both paired-end and singleton reads were mapped to the reference sequences using Minimap2^23^ v2.28-r1209 with the -x sr preset for Illumina reads and the -x map-ont preset for Nanopore reads. Resulting BAM files were merged and sorted using SAMtools^24^ v1.19.2 (merge and sort), with default parameters.. The reference genomes of the species in the ZymoBIOMICS Microbial Community Standard were downloaded from https://zymo-files.s3.amazonaws.com/BioPool/ZymoBIOMICS.STD.refseq.v3.zip. However, for *Lactobacillus fermentum* and *Saccharomyces cerevisiae*, the GenBank assemblies GCF_030770375.1 and GCA_030867715.1 were used, respectively, because the reference genomes provided by Zymo exceeded the expected genome size for these species, likely due to low assembly quality (and, in the case of *S. cerevisiae*, possibly related to ploidy). For the same reason, *Cryptococcus neoformans* was excluded from the analysis, as a high-quality reference genome could not be identified. For *B. dorei* NCBI RefSeq assembly GCF_013009555.1 was used. Genbank accession JQ995537.1 was used as crAssphage reference genome. For the 16S rRNA genes, all bacterial sequences of the SILVA database^25^ (release 138.2) were clustered using MMSeqs2^26^ v18.8cc5c to clusters with 99% identity. The resulting cluster representatives were used as references. The resistance database PanRes^27^ v1.1 and KMA^28^ v1.4.15 with the -1t1 option enabled were used for detection of resistance genes. Which subset of PanRes 1.1 genes were used for the detection of *bla*_CTX-M-1_ and *vanA* genes was based on searching for the qPCR primer sequences, allowing for a 10% mismatch (see Supplementary Table 6). For the ARG trend analysis, only PanRes 1.1 genes that were present in ResFinder were considered.

### Depth estimation and calibration

To account for sequencing variability, depth values and their corresponding error estimates are determined using the following procedure.

MGCalibrator takes a coordinate-sorted BAM file as input and filters reads to retain only those with ≥ *x*% sequence identity to the reference using CoverM^29^ v0.7.0. Optionally, clustering (e.g., grouping alleles of the same gene) and binning (e.g., aggregating contigs belonging to the same MAG) files can be provided to organize reference sequences and their associated reads, using the –cluster and –bin options, respectively. For each reference, a Monte Carlo simulation is performed in which the number of reads mapped to the reference is varied across simulations (see Supplementary Figure 6). Simulated read counts are drawn from a Poisson distribution (λ = number of mapped reads). For each simulation, the depth of coverage is calculated as the mean of the middle *x*% of values. The final depth (*D*_0_) and associated asymmetric uncertainty (Δ*D* _*up*_ and Δ*D* _*down*_) are determined as the mean and the 2.5th and 97.5th percentiles of all simulations.

To account for variation in DNA quantification, these depth values are calibrated using a scaling factor (*k*), calculated as the ratio of DNA amount in the eluate (derived from a Qubit measurement) to the sequenced DNA weight. The scaling factor carries a symmetric uncertainty (Δ*k*) based on DNA weight measurement error. Calibrated depth values are obtained by multiplying the original depth by the scaling factor as seen in Eq. (1).

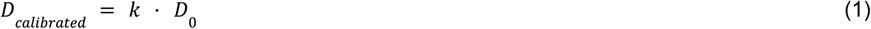

The associated upward and downward uncertainties propagate according to Eqs. (2) and (3), respectively.

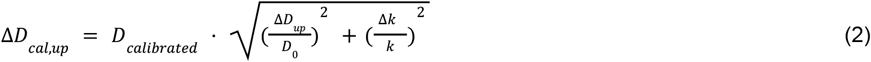

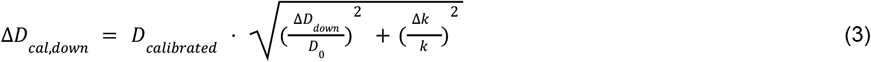

This approach yields calibrated gene or genome copy numbers with propagated errors.

### MGCalibrator settings

The following settings were used for MGCalibrator. For the genes we applied the clustering option, for fragmented genomes in the synthetic dataset and the mock community we used the binning option. All analyses were done with 200 Monte Carlo simulations and using the interquartile mean (i.e., the mean of the middle 50% of values) for calculating depth. A conservative relative Qubit measurement error of 0.0569 was used, estimated from manufacturer-reported performance characteristics, including deviation (accuracy) and coefficient of variation (precision) (Thermo Fisher Scientific^30^). For *B. dorei* and crAssphage we applied read filtering of 97% identity. For the 16S rRNA genes, no read-level filtering was applied because substantial biological variability is expected across 16S sequences, and our aim was to capture the full diversity of 16S gene variants. For the ARGs *bla*_CTX-M-1_ and *vanA*, filtering was also omitted, as KMA inherently produces high-identity mappings, in contrast to the lower baseline identity often observed with Minimap2. For the Zymo microbial community whole-cell and DNA standard samples, 100 Monte Carlo simulations were performed, and 97% and 80% identity filtering was used for Illumina and Nanopore datasets, respectively.

### Synthetic microbiome dataset creation

The synthetic sequencing reads were generated using MAGICIAN^31^ v1.1.2, executing only the Snakemake rules up to the CAMISIM^32^ v1.2 with the included, updated error profile.The MBARC-26^33^ reference genomes formed the base of the dataset, composed of 26 strains (23 bacterial + 3 archaeal) from diverse taxonomic groups, spanning 10 phyla and 14 classes. Four spike-in genomes were added: Influenza A (GCF_000865725.1), CrAssphage (JQ995537.1), *Vibrio cholerae* (GCA_021431945.1), and *Edwardsiella ictaluri* (GCA_042638385.1).

We simulated community composition in two steps. First, chromosome relative abundances were drawn from a log-normal distribution (*meanlog = 1, sdlog = 1*.*5*). Plasmids were modeled as separate “genomes” with abundances equal to its host‐genome abundance multiplied by a copy number. Copy numbers were sampled from truncated normal distributions reflecting their size—small (<10 kb) at 10-25 copies, medium (10-100 kb) at 5-10 copies, and large (>100 kb) at 1-5 copies. The resulting community composition was then used to simulate 5 Gb, 0.5 Gb, and 0.05 Gb of sequencing data, allowing us to evaluate the tool across a wide range of sequencing depths (Supplementary Table 1).

### ARG trend analysis

Phenotype and resistance class abundances were computed by summing the abundances of all annotated genes, with upper and lower uncertainties propagated separately in quadrature. Temporal trends in gene abundances, phenotypes, and resistance classes were evaluated per city and dataset by retaining features with sufficient coverage (depth >10 for ≥5 features in Utrecht and ≥6 features in Houtrust). Temporal trends were determined using the non-parametric Kendall **τ** test. For clarity in visualization, only the five features with the largest absolute Kendall **τ** per dataset and city were plotted, with error bars reflecting the original measurement uncertainty.

## Data Availability

All the sequence reads have been deposited at the European Nucleotide Archive (Bioproject PRJEB107319) and the individual metagenomic read accessions, along with measured DNA weights, can be found in Supplementary Table 3. The MGCalibrator software can be found at github.com/NimroddeWit/MGCalibrator. A Docker image (version 1.0.0) is available on Docker Hub (https://hub.docker.com/r/amulyabaral/mgcalibrator) and archived on Zenodo (DOI: 10.5281/zenodo.18773041). qPCR measurements can be found in Supplementary Table 7.

## Acknowledgements

We thank the Novo Nordisk Foundation (Grant: NNF24SA0094147), the Dutch ministry of Health, Welfare and Sport, and the Dutch ministry of Agriculture, Fisheries, Food Security and Nature for supporting this work.

