## Supplementary Information for "Calibrating for absolute microbiome abundances without spike-ins"

### Supplementary Figures

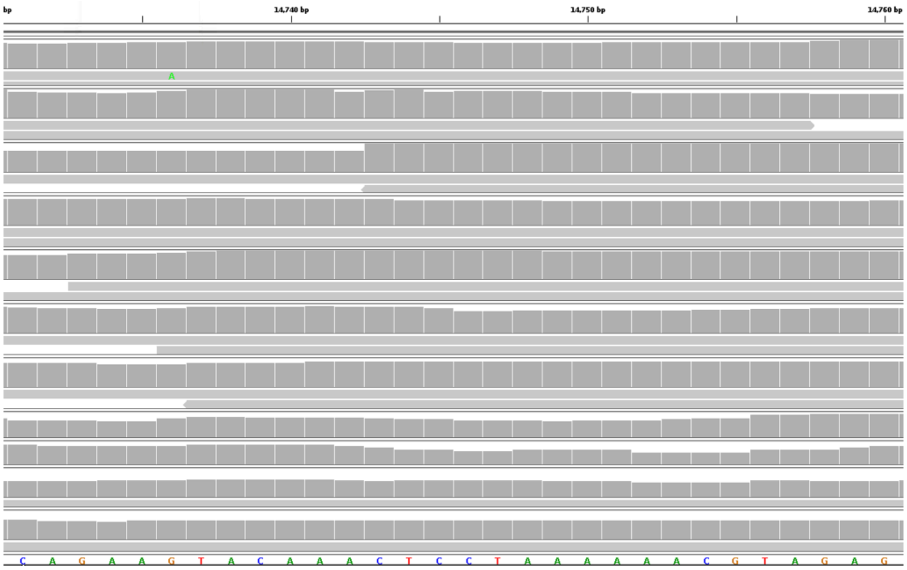

b

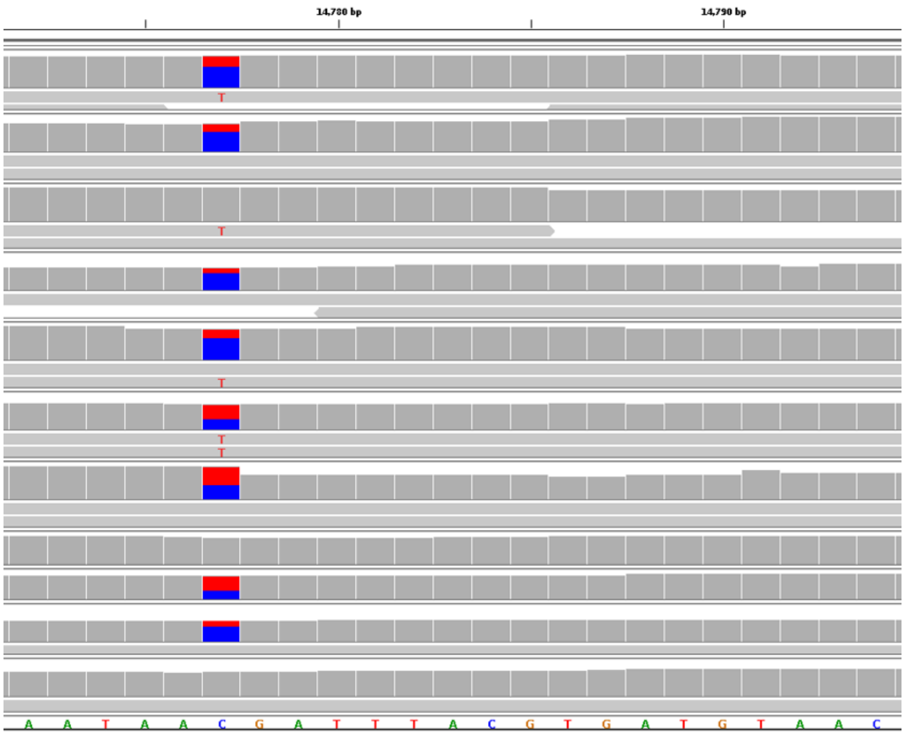


c

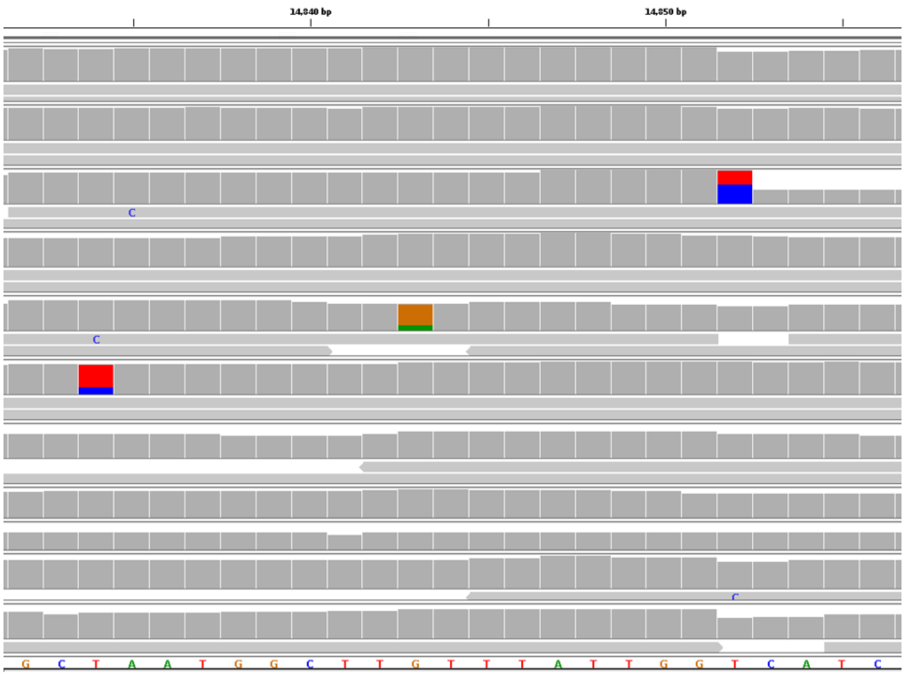


***Supplementary Figure 1. Mapped reads at qPCR primer sites are highly consistent with the crAssphage reference genome.*** *For a random subset of 11 datasets, the alignment of reads against the primer sites (a: forward, b: probe, c: reverse) was examined to assess sequence matching with the crAssphage reference genome. Perfect alignment was observed for the forward primer. A subset of the sequenced reads exhibited a single nucleotide mutation*

*(C>T) relative to the probe sequence, while sporadic mutations were detected in the region corresponding to the reverse primer.*

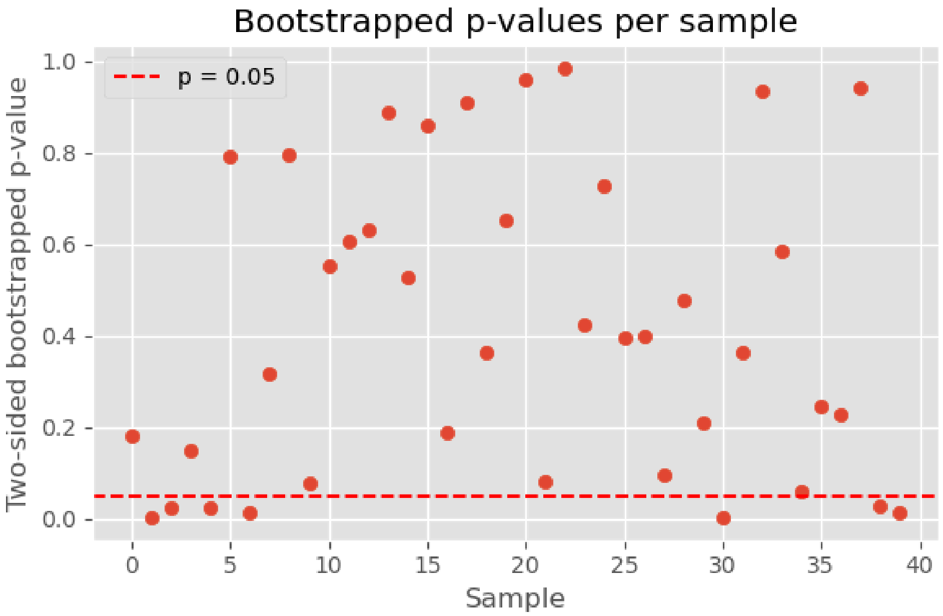


***Supplementary Figure 2. Depth at crAssphage qPCR site is not deviating from the depth across the whole genome.*** *To test whether the sequencing depth at the qPCR target site was representative of the overall genome coverage for crAssphage, we performed a bootstrapping analysis. For each dataset, the mean coverage of the 126 bp qPCR region was compared to the distribution of mean coverages of randomly sampled contiguous regions of the same length across the whole genome. In most datasets, the two-sided bootstrapped p-values were above the conventional significance threshold ($p > 0.05$), indicating no significant difference between the qPCR site and the genome as a whole. (Only a small number of datasets showed significant deviation (two-sided $p < 0.05$), suggesting possible local amplification or sequencing biases in these cases.)*

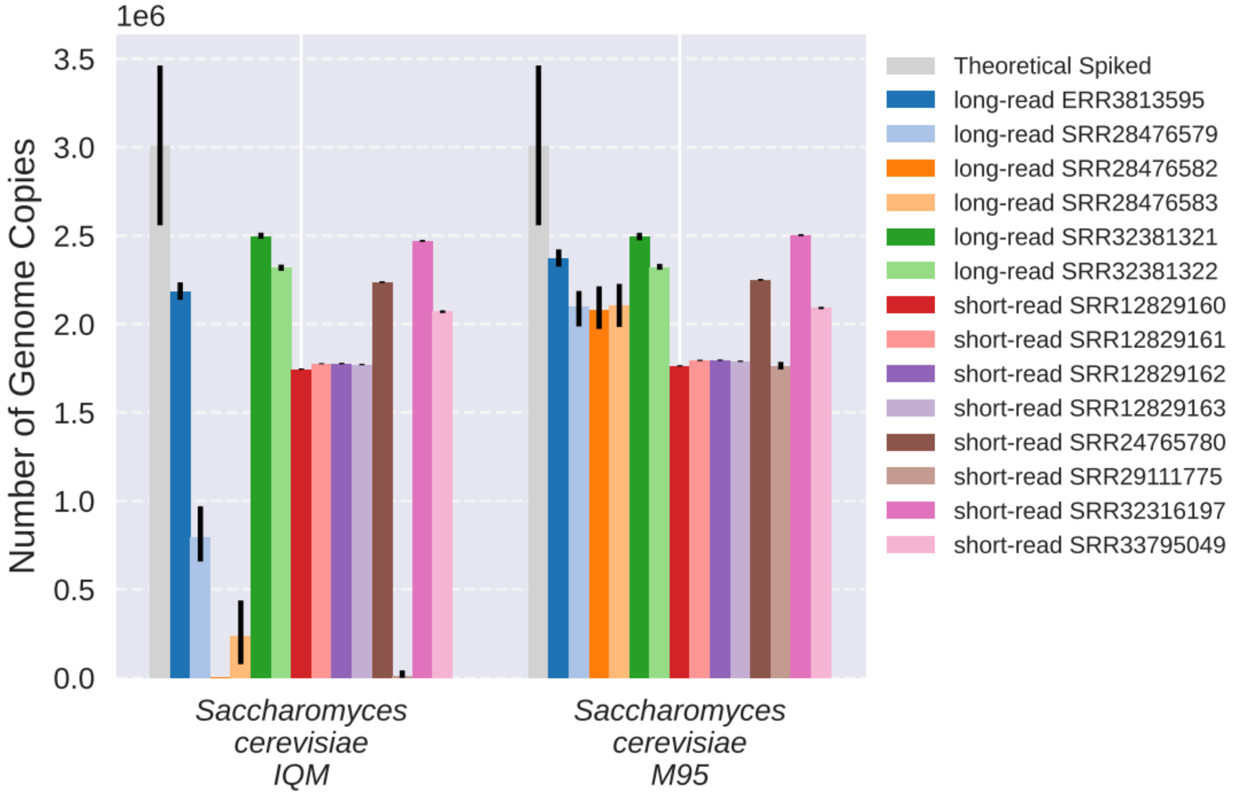


***Supplementary Figure 3. Comparing IQM and M95 depth calculation methods for calibration of*** **Saccharomyces cerevisiae*.***

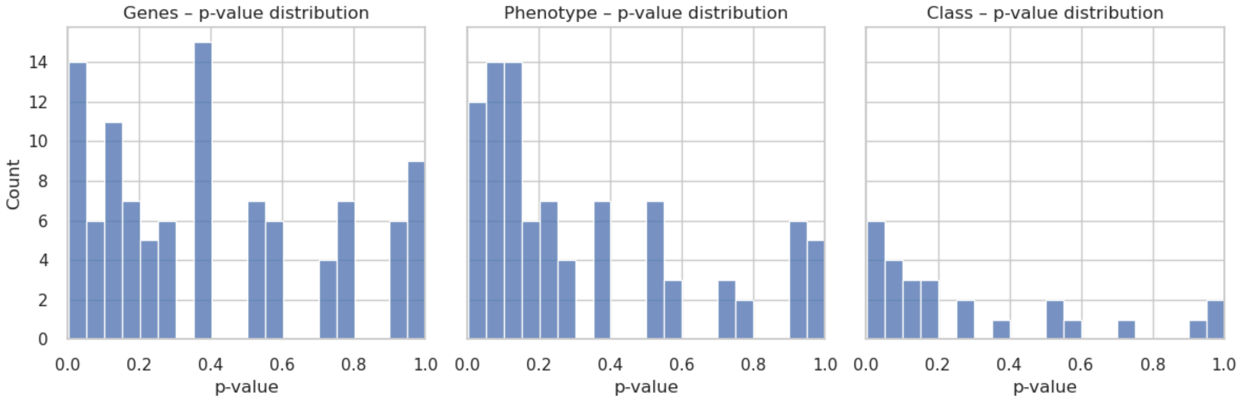


***Supplementary Figure 4. Kendall τ raw p-values of resistance genes, phenotypes and classes.***

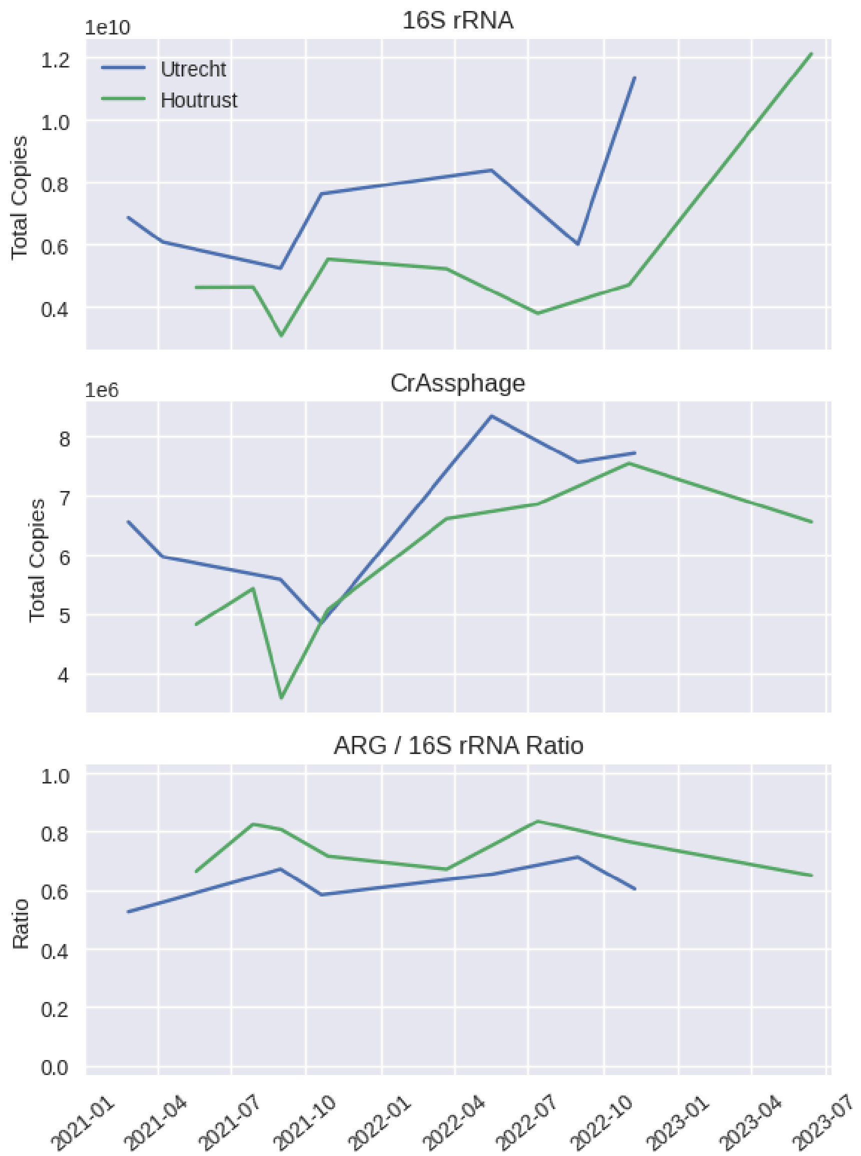


***Supplementary Figure 5. Temporal trends in total 16S rRNA gene copies, total crAssphage copies, and the ARG/16S rRNA gene ratio at the Utrecht and Houtrust sampling locations.*** *Total copy numbers were estimated using MGCalibrator. Total ARG depth was derived from KMA output by summing all values in the* Depth *column.*

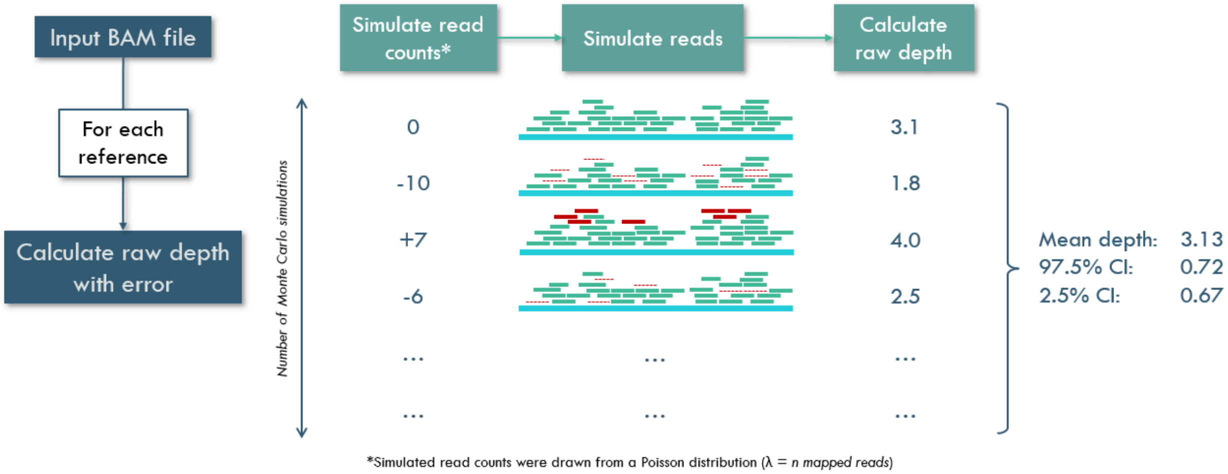


***Supplementary Figure 6. Schematic overview of depth calculation with error estimation using Monte Carlo simulation****. A BAM file is used as input. For each reference, the depth and corresponding error estimates are calculated. In each Monte Carlo simulation, a read count is drawn from a Poisson distribution (λ = number of mapped reads), and reads are removed from or added to the original set accordingly. Depth is calculated for each simulation. Finally, the mean depth across all simulations is computed together with the corresponding 95% confidence intervals.*

### Supplementary Tables

***Supplementary Table 1. Species composition of the synthetic microbiome samples.*** *For each species the relative abundance and the theoretical abundances at three sequencing depths (5 Gb, 0.5 Gb, 0.05 Gb) are shown.*

***Supplementary Table 2. Zymo Microbial Community DNA Standard datasetssamples downloaded from the European Nucleotide Archive (ENA).*** *Both Illumina and Nanopore datasets were downloaded from ENA (https://www.ebi.ac.uk/ena/browser/home).*

***Supplementary Table 3. Information on the 42 samples sequenced in this study.*** *DNA concentrations were measured using a Qubit® dsDNA HS Assay Kit, and all samples were sequenced on the Illumina NextSeq 2000 platform. Additional sample data, like the number of raw reads and the scaling factor used by MGCalibrator are also provided.*

***Supplementary Table 4. qPCR information.*** *Information on the five qPCR runs performed for this study, including details on the target genes, primers, and cycling conditions.*

***Supplementary Table 5. Sequences of two in-house designed Gblocks synthesized by Integrated DNA Technologies (IDT).*** *The first Gblock was originally designed to target 8* *different loci, but in this study, it was used for only two targets: crAssphage and* Bacteroides dorei*. The second Gblock was utilized for three targets: the 16S rRNA gene,* vanA*, and* $bla_{CTX\text{-}M}$.

***Supplementary Table 6. PanRes genes for* vanA *and* $bla_{CTX\text{-}M}$.** *A list of all PanRes genes used for the detection of antibiotic resistance genes* vanA *and* $bla_{CTX\text{-}M}$. *Relevant PanRes genes were identified by searching for primer sequences specific to each resistance gene, allowing a 10% mismatch to account for sequence variability.*

***Supplementary Table 7. Number of gene copies as detected by qPCR for the 16S rRNA gene,* Bacteroides dorei *(HF183), crAssphage,* $bla_{CTX\text{-}M}$*, and* vanA.**
